# Translational landscape during Seed Germination at Sub-Codon resolution

**DOI:** 10.1101/2025.06.24.661055

**Authors:** Bing Bai, Run Qi, Wei Song, Harm Nijveen, Leónie Bentsink

## Abstract

Seed germination is crucial for agricultural reproduction. A deep understanding of this process can secure healthy growth at the early phases of plant development and therefore the yield. Recent research indicates that germination is a complex process involving translational regulation. A large group of seed stored mRNAs together with newly synthesized transcripts are regulated by post-transcriptional mechanisms and selectively translated at different stages to support the germination process. To investigate the mechanism of translational control, we performed ribosome profiling on mRNAs of distinct physiological stages during *Arabidopsis thaliana* seed germination. The 3-nucleotide periodicity of the ribosome association on mRNAs identified indicates their translational potential. Dry seeds, in which translation is on hold are characterized by a unique ribosome association landscape with high association at the 5’ and 3’ UTR, compared to physiological stages that show active translation. Start codon specific stalling of ribosomes in dry seeds is associated with an adenine enriched sequence motif. Throughout germination codons encoding glycine, aspartate, tyrosine and proline are the most frequent ribosome pausing sites. Moreover, the non-coding RNAs which we identified as associated to ribosomes are indeed translated as was revealed by investigating total seed proteome data. Seed specific upstream Open Reading Frames (uORFs) are identified which may be associated to translational control of adaptative response during early seed germination. Altogether, we present a first ribosome profiling analysis across seed germination that illuminates various regulatory mechanisms that potentially contribute to the seeds survival strategy.

## Introduction

Seeds are the primary structures responsible for plant propagation and the survival strategy of plant. During germination, the seed embryo develops into a seedling, which subsequently grows into a mature plant. In their fully matured state, seeds store large amounts of mRNA. The translation of these stored mRNAs is essential for germination to take place (Rajjou et al., 2004). These stored mRNAs can survive for years, while still maintaining their capacity for translation (Dure and Waters, 1965; Waters and Dure, 1966; Yashina et al., 2012; Sano et al., 2016). In *Arabidopsis thaliana*, stored mRNAs are predominantly associated with monosomes, single ribosome units, that are not actively involved in translation (Bai et al., 2020). When dry seeds take up water, translation of monosome-associated transcripts is initiated. As more ribosomes bind to a single mRNA molecule, a polysome is formed, allowing multiple proteins to be synthesized simultaneously from a single mRNA. This process is strongly regulated and referred to as the Hydration Translational Shift (from dry seeds until 6 hours after seed imbibition). A second translational shift occurs around the moment that the seed embryonic root (radicle) breaks through its surrounding layers, the testa and the endosperm, and is referred to as the Germination Translational Shift (Bai et al., 2017). The translational shifts referred to indicate a change in translational efficiency (TE), e.g. the proportion of ribosome bound mRNA over total mRNA, a ratio used to evaluate the extent of translation for specific mRNAs (Ingolia et al., 2009; Liu et al., 2012; Juntawong et al., 2014).

The fate (synthesis, translation or degradation) of ribosomes on a transcript is influenced by interactions between the ribosome and mRNA sequence features such as the 5’ Untranslated Region (5’ UTR), upstream Open Reading Frames (uORFs) and the 3 ‘Untranslated Region (3’ UTR). Therefore, accurately determining the position of the ribosome on stored mRNAs in seeds can provide insights into the state of mRNA storage and their fate during seed germination when the translation is re-activated. Information about the ribosome position on the mRNA can be revealed by ribosome profiling. Deep sequencing of ribosome protected mRNA fragments (ribosome footprint, RP) identifies the position of the ribosome on the RNA molecules. This technique has been extensively used across various species, including yeast, bacteria, Archaea, Drosophila, and plants, to investigate translational control in response to environmental stimuli and during developmental processes (Ingolia et al., 2009; Hsu et al., 2016; Zhang et al., 2017; Gelsinger et al., 2020). In crop seeds ribosome profiling has been used to study translational control during seed maturation (Guo et al., 2023) and to study tissue specific translation (Zhu et al., 2023).

Sequence features such as the length of the UTRs, GC content and RNA structure are associated with mRNA translational efficiency in Arabidopsis (Bai et al., 2017). Moreover, uORFs that are located in the 5’ UTR of some eukaryotic mRNAs, upstream of the main ORF (mORF) are known to regulate translation. These uORFs can cause stalling or releasing of the ribosome from the mRNA and thereby inhibit translation of the mORF (Meijer and Thomas, 2003; Wiese et al., 2004). The uORFs translational control mechanism is conserved, across multiple plant species the interaction between the uORF encoded peptides and metabolites, such as sucrose, polyamine, thermospermine and other metabolites, have been identified (van der Horst et al., 2020).

In addition to mRNAs, eukaryotic cells contain a substantial number of non-protein coding RNAs. Among these, long non-coding RNAs (lncRNAs) are defined a transcript longer than 200 nt that either lack an ORF entirely or do not contain an ORF encoding more than 100 amino acids. LncRNAs are known to play important regulatory roles in plants. For example some have been shown to encode functional peptides that are associated with plant morphogenesis and stress-induced plasticity (Bazin et al., 2017; Zeng et al., 2018; Wang et al., 2019; Yu et al., 2019). The majority of lncRNAs are transcribed by RNA Polymerase II and contain typical mRNA-like features, such as 5’m7G cap and a 3’poly (A) tail. Therefore, lncRNAs are processed in the same way as mRNAs (Wu et al., 2017). LncRNAs are characterized by a lower conservation and abundance, more tissue-specific expression, and lower splicing efficiency than mRNAs (Ulitsky and Bartel, 2013). Over a thousand lncRNA are identified as associated with ribosome which indicates they have the potential to be translated (Bazin et al., 2017; Zeng et al., 2018).

To obtain a better understanding about translational regulation during seed germination we performed ribosome profiling (ribosome footprinting, Ribo-seq). We characterize the features and dynamics of the ribosome associated mRNAs in both dry seeds and during seed germination. Moreover, we utilize transcriptome-wide ribosome position maps across the course of seed germination to explore the function of uORF mediated translational control and to identify lncRNAs with a translation potential.

## Materials and Methods

### Seed Material Preparations

Seeds of the *Arabidopsis thaliana* accession Columbia-0 (NASC N60000) were used for ribosome profiling. Sampling was performed according to Bai et al (2017). Briefly, seeds were germinated in a 22°C incubator under continuous light (143 μmol m^-2^s^-1^). Seed were harvested at 0, 6, 26, 46 and 72 hours after imbibition (HAI). The harvested material was frozen in liquid nitrogen followed by freeze-drying. The dry material was stored at -80°C until further analysed.

### Ribosome isolation and footprint preparation

The ribosome isolation was performed according to Bai et al (2017) and the footprint preparation was modified from Hsu et al (2016). In brief, 50 mg dry material was extracted with 2 ml of polysome extraction buffer, PEB (400 mM Tris pH 9.0, 0.25M sucrose, 200 mM KCl, 35 mM MgCl_2_, 5 mM EGTA, 1 mM phenylmethane sulfonyl fluoride, 5 mM Dithiothreitol, 50 μg/mL Cycloheximide, 50 μg/mL Chloramphenicol). A 50-μL aliquot of supernatant was used to extract total RNA for constructing RNA-seq libraries (see below). A 200-μL aliquot of supernatant was used for ribosome footprint preparation. This aliquot was treated with RNase I nuclease (Invitrogen, cat. no. AM2294) at 10 U per 10 OD 260 unit ribosome aliquot. Enzyme digestion was performed at 23 °C with gentle mixing on a thermal mixer (Eppendorf) for 1.5 hour. Nuclease digestion was stopped by adding 15 μL of SUPERase-in (Thermo Fisher Scientific; AM2696). Fragmented RNA was purified and separated by 15% (wt/vol) TBE-urea PAGE (Thermo Fisher Scientific; EC68852BOX), and gel slices corresponding to fragment sizes between 26 and 34 nt were excised according to the two RNA oligo markers with 26 and 34 nt in size. Ribosome footprints were recovered from the excised gel slices following the overnight elution method as specified by Ingolia et al (2012).

### Ribosome and RNA sequencing library construction and sequencing

After obtaining ribosome footprints above, Ribo-seq libraries were constructed according to ARTseq/TruSeq Ribo Profile Kit manual and amplified by 13 cycles of PCR with a barcode incorporated in the primer. The PCR products were gel purified using the overnight method described by Ingolia et al (2012b). Equal molarities of the libraries were pooled for single-end 50-bp sequencing using an Illumina NovaSeq 6000 sequencer.

For RNA-seq, a 50-μL aliquot of supernatant as described above was used to extract total RNA by TriPure Isolation Reagent (Roche, Basel, Switzerland), followed by clean-up using RNeasy Mini spin columns (Qiagen, Hilden, Germany). The concentration of purified RNA was quantified using nanodrop. Then, 5 μg of total RNA was subjected to rRNA depletion using Illumina Ribo-Zero Plus rRNA Depletion Kit following the manufacturer’s instructions. The rRNA-depleted RNA was used to construct sequencing libraries using the ARTseq/TruSeq Ribo Profile Kit (Illumina). The circularized cDNA was amplified by 11 cycles of PCR and gel purified using the same procedure for the Ribo-seq libraries described above. Libraries were barcoded, pooled, and sequenced using an Illumina HiSeq 2500 machine. The sequencing data were submitted to Sequence Read Archive (SRA) from NCBI with BioProject ID PRJNA1169673.

### Adaptor Processing, Quality Assessment, and Alignment

FASTQ files were obtained and all data analysis steps were performed in house, using a combination of command line software tools and python scripts. Adaptors were trimmed off with the cutadapt program (Martin, 2011) using the following adaptor sequence: CTGTAGGCACCATCAAT, reads smaller than 18 bp were removed; Quality reports of raw and trimmed read sets were generated with the FastQC program. The getfasta command from the bedtools program (Quinlan and Hall, 2010) was used to extract strand-specific sequences in FASTA format for the annotated rRNA, tRNA, miRNA, snRNA, snoRNA and lncRNA sequences in the Arabidopsis genome using the Araport11 genome annotation (Cheng et al., 2017). The trimmed and quality-filtered reads were mapped to these RNA sequences with the short sequence sensitive mapping tool Bowtie v1.2.3, following the strategy of mapping, filtering unmapped reads and mapping to another reference in the reference order of rRNA, tRNA, snRNA, snoRNA, miRNA, lncRNA, mRNA and genome (Langmead et al., 2009). Each read was only allowed to map once to the best location with at most one mismatch and only to the coding strand except in the final step when mapping to the genome. The Arabidopsis genome sequence was retrieved from The Arabidopsis Information Resource (https://www.arabidopsis.org), version 10 and the Araport11 genome annotation was download from Ensembl Plants (https://plants.ensembl.org).

### Metagene Analysis, Data Visualization

In order to verify and select specific reads showing periodicity of ribosome association, the R package RiboWaltz was used (Lauria et al., 2018). Transcriptome alignment (binary alignment map, BAM) files and Araport11 annotation gff file were provided as input (Cheng et al., 2017). For data visualization, the corresponding FASTA sequences (e.g. lncRNA, transcriptome, genome) generated for Bowtie mapping were used as reference. BAM files were filtered for specific ribosome protected read length and visualized using IGV genome browser (Robinson et al., 2011).

### Differential Translation Comparison, Clustering, and Gene Ontology Term Enrichment Analysis

Raw read counts were normalized with the DESeq2 R package and PCA analysis was performed using the plotPCA function from DESeq2 based on normalized read counts. Differential translation analysis was performed by counting reads in the transcriptome alignments using the featureCounts function from the Subread v 2.0.1 R package with gff files as reference (Liao et al., 2013). DESeq2 was applied to perform differential translation analysis between 5 time points. Statistical analysis was performed using the design formula ∼ assay + condition + assay:condition. Using the likelihood ratio test of DESeq2, which removes the interaction term in the reduced model, the difference in translational efficiency by comparing the ribosome-associated RNA to total RNA between two conditions can be tested. The length normalized counts in rpkm for each gene were calculated based on the formula rpkm = CDS mapped reads / (CDS length /1,000 * total reads/1,000,000) by using normalized read counts. Genes with an 5% FDR adjusted *p*-value < 0.05 were considered as translationally regulated. Read counts of various feature types of interest (e.g. CDSs, UTRs) were calculated separately using the featureCounts function from Subread v 2.0.1 with gff files as reference. For Gene Ontology (GO) enrichment Fisher’s exact test was performed using the SciPy Python library (version 1.5.2) and a 5% FDR was applied for multiple testing using the multitest.multipletests function with method Benjamini/Hochberg from the Python statsmodels module (version 0.12.1). As GO annotation the GO_slim file from the TAIR website (https://www.arabidopsis.org/) was used. The start codon-enriched genes in dry seeds were identified by calculating the proportion of P-sites at every position across the entire CDS. Genes with a P-site proportion at the start codon exceeding 10% of the total P-sites were selected and with a start codon P-site count exceeding 30 were further retained as start codon-enriched genes. The PANTHER server (https://pantherdb.org) was used for GO enrichment analysis (release 20221013) (Thomas et al., 2022).

### MEME motif analysis and enrichment

Motif enrichment analysis was performed on RNA sequences of selected genes using MEME (version 5.5.5) to identify significant motifs (Bailey et al., 2015). The search was conducted with the “One Occurrence Per Sequence” (OOPS) model, setting a maximum of 3 motifs with widths between 6 and 10 nucleotides for the 176 translation initiation enriched genes. Background dinucleotide frequencies were provided separately for the defined start codon context region of the transcript all genes detected in the RNA-seq. To test the specificity of the resulting motifs, FIMO (Bailey et al., 2009) was used to scan all genes detected in the RNA-seq in the defined start codon context region. Motifs with a *p*-value ≤ 0.001 were considered significant hits. Obtained motif counts were used to compute the enrichment *p*-value for the gene lists versus the background by means of a one-tailed Fisher’s exact test, performed with a custom script written in R (http://www.r-project.org/).

### Long non-coding RNA analysis

Long non-coding RNA in both total RNA (total lncRNA) and ribosome associated RNA (ribo lncRNA) are classified with the Araport11_gene_type file (2019 update) from the Araport11 genome release. lncRNA with rpkm > 5 in all three biological replicates were kept for both total lncRNA and ribo lncRNA. Translational efficiency (TE) of lncRNA was calculated in the same way as for mRNA, as the ratio of ribosome-bound lncRNA to total lncRNA. lncRNA that were detected both in RNA-seq and Ribo-seq and lncRNAs with detected ORFs with predicted ORFs by systemPipeR that were significantly changed in TE were considered “translationally regulated”.

### Mass-spectra based proteomic analysis of lncRNA encoded peptides during seed germination

The peptide sequence encoded by the short Open Reading Frame (sORF) of the identified lncRNAs was predicted by systemPipeR’s predORF function (TW and Girke, 2016) and stored in FASTA format as input for proteomic analyses. The proteomic data was retrieved from Bai et al. (2021) at ProteomeXchange with accession PXD027345 for mass spectrometry based proteomic analysis during seed germination. Identification of proteins from the raw MS Data was done using Proteome Discoverer (PD) 2.4 SP1 (Thermo Fisher Scientific, USA) with default settings. PD was set up to search the protein database of *Arabidopsis thaliana*, downloaded from TAIR (https://www.arabidopsis.org/, version TAIR10_pep_20101214). Human keratin and protease mixture sequences were used as the contaminant database for proteomic searching. The processed data output was then exported to Excel for further usage. The peptides identified from all biological replicates in each stage were selected as the identified peptides encoded by the identified lncRNAs.

### uORF analysis

SystemPipeR’s predORF function was used for uORF prediction by searching potential ORFs on 5’ UTR region of each gene transcript. ORFs longer than 60 bp (20 aa) were kept and converted into uORF annotation in gff format. 5’UTR sequences were extracted based on the Araport11 genome and annotation. Ribosome associated uORFs and ribosome associated CDSs reads were extracted from Ribo-seq data based on each annotation respectively. Genes with both ribosome-associated uORFs and CDSs with rpkm > 5 in all three biological replicates were kept for differential expression (ribosome abundance) analysis. Ribosome associated uORFs and ribosome associated CDSs were calculated separately and compared with each other. Comparisons between CDS and their associated uORFs were calculated using the generalized linear model of DESeq2 as described above, using read counts from Ribo-seq of CDS and uORF as input to DESeq2. Genes with significant changes in ribosome footprint (RF) count in upstream open reading frames (uORFs) and coding sequences (CDS) were identified using DESeq2, with adjusted *p*-values calculated using the Benjamini–Hochberg method (padj < 0.05). Translationally regulated genes were defined as those showing opposite regulation patterns in log_2_Fold change in RF count of uORF and CDS between consecutive time points during seed germination.

## Results

### Ribosome profiling during seed germination

To determine the precise position of ribosomes on the mRNAs during translation, ribosomes were isolated during seed germination at 0, 6, 26, 48 and 72 hours after imbibition (HAI). These time-points correspond to the physiological stages: dry seeds, seeds at early imbibition, seeds before testa rupture, seeds at 80% of endosperm rupture and 80% of seedling greening. These stages were shown to exhibit extensive translation dynamics (Bai et al, 2017). Transcriptome-wide ribosome profiling (Ribo-seq) was performed at a depth of ∼100 million reads per sample. Reads mapping to rRNA and tRNA, and unmapped reads were removed. Reads mapping to non-coding RNAs were stored as a separate dataset for further analyses (Supplemental Table 1). About 10% of the total number of reads per sample mapped to an mRNA (Supplemental Table 1). Periodic patterns in the RNA sequence strongly indicate ribosome attachment, as ribosomes advance along the RNA in increments corresponding to three nucleotides (codons). A strong 3-nt periodicity was found for reads or ribosome footprints (RF) in the size range between 27-nt and 29-nt. In dry, 6 and 26 HAI seeds the 28-nt RFs are most pronounced, whereas at the later time-points during seed germination the 29-nt RFs become more pronounced (Figure 1a, Figure S1). The -12-nt and +15-nt position marked the ribosome extremities at the 5’ and 3’ respectively at the start codon context (Figure 1a). At the stop codon, in addition to the 3-nt periodicity we also observe footprints in the +1 reading frame, both at the 5’ and 3’ ribosome extremities (Figure 1a and b), suggesting a ribosomal frameshift. The ribosome P-site serves as a reliable reference point for ribosome positioning, since the start codon is aligned with the P-site during translation initiation. In order to further validate the genuine RFs, the P-site position was visualized across the up- and down-stream sequences of start and stop codon context. Like the ribosome extremity plot, the ribosome P-site metaplot also shows a clear periodic distribution (Figure 1b). This periodic P-site signal was identified in the CDS at all physiological stages including a strong general peak at the start codon and a smaller peak at the 5^th^ codon position (Figure 1a,b Supplemental Table 2), indicative of initiation and post-initiation translational pausing (Han et al., 2014). As expected, in dry seeds the peak signal was overall lower compared to the imbibed stages, (Figure 1a and b, Supplemental Table 2), consistent with the repressed translational activity in dry seeds. Strikingly, in dry seeds ∼7% of the total mapped reads were positioned at start codon, this corresponds to 40% of the reads that are positioned at the P-site of the ribosome in the displayed region of the metaprofile (Figure 1b). This is significantly higher than in the other stages (∼1%-2% for 6-72 HAI; t-test, *p*-value < 0.05, Supplemental Table 3), corresponding to 10-20% of the reads that are positioned at the P-site of the ribosome (Figure 1b). Summing up all the RFs across the transcripts, 92.3% of the RFs in dry seeds mapped to protein-coding sequences (CDSs), 3.57% to 5′ UTRs, and 4.13% to 3′ UTRs, while at other time points ∼96%-98% of the RFs mapped to CDSs, ∼1.9-2.3% to 5′ UTRs, and ∼0.8%-1% 3′ UTRs (Figure 1c, Supplemental Table 2, 3), indicating a reduced ribosome loading in CDS and an increased loading in both 5’ and 3’ UTRs in dry seeds compared to other physiological stages (*p*-value <0.05).

**Figure 1.**
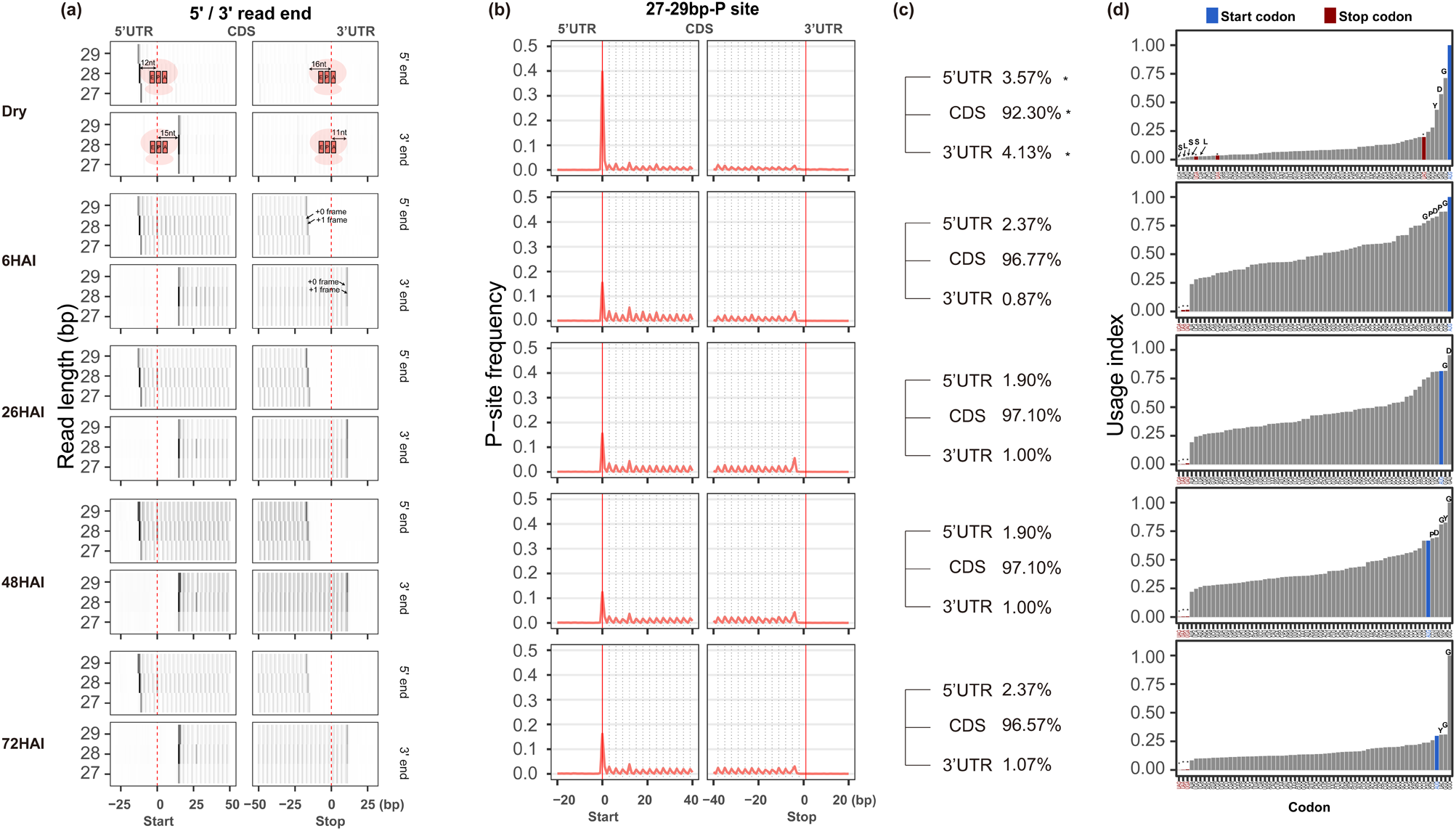
Ribosome dynamics during seed germination. (a) Meta-gene heatmap of ribosome footprinting around start and stop codons. Heatmaps display aggregate ribosome occupancy based on the 5’ end (top panels) and 3’ end (bottom panels) of ribosome-protected fragments (RFs), aligned relative to annotated start and stop codons. Signals are shown for RFs of 27, 28 and 29 nt in length across five stages of germination: dry seeds and seeds at 6, 26, 48 and 72 hours after imbibition (HAI). For the dry seed panels, a schematic ribosome is superimposed on the plots. The E, P and A site position and the distance of the P-site to the start and stop codon are indicated to illustrate ribosome positioning (b) Metaprofile of ribosome occupancy displaying the relative abundance of P-sites around the start and the stop codon of annotated CDSs (fragment lengths, 27-29 nt combined) during seed germination. (c) The percentage of RFs in the 5’ UTR, CDS and 3’ UTR at each stage during seed germination. Asterisk indicate the significance comparing percentages of in dry seeds compared to all other stages (*p*-value < 0.05) (d) Codon usage based on in-frame P-sites during seed germination. The codon usage index is calculated as the frequency of in-frame P-site positions associated with each codon along CDSs, normalized for codon frequency. For the most frequent codons, their corresponding amino acid is displayed above each bar. The blue and red bars represent the start and stop codons respectively. Note, the AUG codon includes both start codons and internal methionine codons within the CDS.

The strength of ribosome association can be influenced by the codons encountered during translation. To identify the codon that contributes the most to ribosome pausing (temporary halting of the ribosome during the process of translation), the codon usage index was computed based on the in-frame P-site position (Figure 1d). In dry seeds, ribosomes were paused mainly at the start codon followed by codons encoding glycine (G), aspartate (D) and tyrosine (Y). Interestingly, ribosomes in dry seeds were also partially paused at stop codons, especially at the UAG codon (codon usage index=0.13, Supplemental Table 4), which might reflect an incomplete translation termination. At 6 HAI, ribosomes were associated most frequently with the start codon followed by codons for glycine (G), proline (P) and aspartate (D), while the ribosome occupancy at the stop codon was strongly reduced at this stage. From 26 HAI onwards, as germination progressed and translation become fully activate, the start codon was no longer the primary ribosome pausing site. Instead, codons for glycine (G), aspartate (D), tyrosine (Y) and proline (P) emerged as the most frequent codon pausing positions. Although no stalling at the stop codon was identified, likely because only the P-site position was analysed, a distinct peak was observed 4 nucleotides upstream of the stop codon (Figure 1b). This feature has been reported previously and likely corresponds to RNA compaction during translation termination (Brown et al., 2015; Heyer and Moore, 2016).

### Translational features of start codon enriched genes in dry seeds

Dry seeds show a dominant RF peak at the start codon (Figure 1a-c), which maps to 176 genes with enriched ribosome occupancy at their start codons (Supplemental Table 5). Gene Ontology analyses showed that these genes are enriched for biological functions with terms as seed oil body biosynthesis, lipid storage and seed maturation (Supplemental Table 6). When investigating the translational behaviour of these genes, two distinct groups were identified. The first group consists of genes that are enriched for RFs at the start codon in dry seeds, but show a marked depletion at 6 HAI. Examples include genes encoding oil body-associated proteins and Late Embryogenesis Abundant (LEA) proteins. Their transcripts are known to accumulate during maturation and are likely degraded immediately upon imbibition, consistent with their reduced transcript level during germination (Figure 2a-b). The second group includes genes that also show RF enrichment at the start codon in dry seeds, but exhibit increased ribosome occupancy in their CDS upon imbibition (6 HAI). Their transcript abundance reduces sharply upon further imbibition (Figure 2c). These genes include seed storage protein genes. The accumulation of seed storage proteins during seed maturation indicates that the translation of these transcripts starts already during seed maturation and continues during early imbibition. These two modes of ribosome association pattern may represent different fates for seed stored mRNAs with varying function during seed germination.

**Figure 2.**
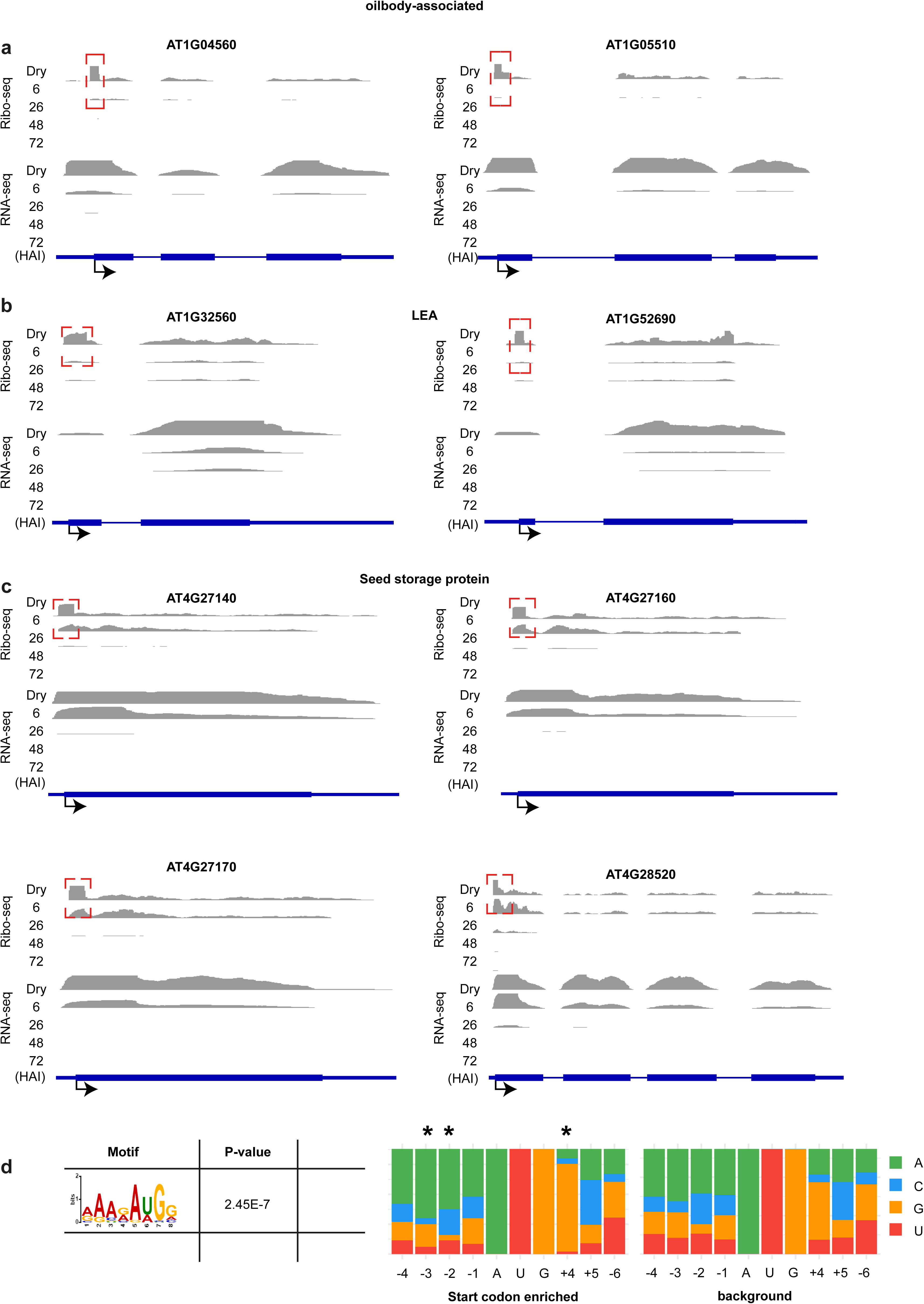
Start codon-enriched ribosome footprints in dry seeds. (a-c) Coverage plots showing translational (Ribo-seq) and transcriptional (RNA-seq) read densities during seed germination for representative genes with start codon-enriched ribosome footprints in dry seeds and at 6 HAI. Genes are grouped by function: (a) oil body associated, (b) Late Embryo Abundant (LEA) and (c) Seed storage protein genes. (d) Left: enriched motif found around the start codon of genes with start codon-enriched ribosome footprints in dry seeds. Right: distribution of each nucleotide per position relative to the start of the CDS in the start codon-enriched RF genes compared to the background genes are shown. The asterisk indicates a significant difference in nucleotide distribution at the -3, -2 and +4 positions (*p*-value < 0.0001, Chi-square test).

To investigate whether there is a sequence basis for ribosome pausing at the start codon, the sequence context surrounding the AUG start codon of the 176 start codon enriched genes was analysed for motif enrichment. This analysis revealed the motif AAAG<u>AUG</u>G (Figure 2d), whose nucleotide composition differs significantly from background sequences at positions -3, -2 and +4 relative to the adenine of the start codon. Specifically, positions -3 and -2 are enriched for adenine, while position +4 shows increased guanine and reduced adenine and uracil (Supplemental Table 7). This motif closely resembles the canonical Kozak consensus sequence in dicot plants AAAMA<u>AUG</u>GC (where M = A or C) (Joshi et al., 1997).

### Translational changes of mRNAs and lncRNAs during seed germination

Earlier, we have evaluated the translational efficiency (TE) of mRNAs during seed germination using polysome profiling (Bai et al., 2017). The shift in TE from dry to 6 HAI seeds that we called the “hydration translational shift” (HTS) is also present in the Ribo-seq results of this study (Figure S2, Supplemental table 8-10). Comparing the differentially expressed genes between consecutive timepoints, we find a large overlap between the RNA-seq used in this study and the previously published microarray results, indicating the robustness of cross-platform translatome profiling during seed germination (Figure S3). At the translational level, the overlap is considerably lower, likely due to methodological differences: the earlier study employed polysome profiling while the Ribo-seq used here also captures monosomes.

The sequencing-based method used in this study allowed us to explore ribosome association with non-coding RNAs such as long non-coding RNAs (lncRNAs) during seed germination. In total, we detected 1594 and 218 non-coding RNAs from the RNA-seq and Ribo-seq datasets based on genome annotation (Supplemental table 11). To further investigate the coding potential of these identified lncRNA in seeds, both Ribo-seq and RNA-seq detected lncRNAs were investigated for the presence of ORFs. These analyses revealed that 463 lncRNAs expressed during seed germination contain a predicted ORF, among which 25 lncRNAs were found to be under translational control at distinct physiological stages (Supplemental table 11 and 12). To validate the translation potential of the ORF containing lncRNAs, the predicted peptide sequences encoded by their ORFs were used for peptide discovery (using Proteome Discoverer (PD) 2.4 SP1 (Thermo Fisher Scientific, USA)). The proteomic data for these analyses was obtained from a total proteome analysis that was performed on the same germination time-points as those analysed in the present study (Bai et al., 2021). In total, 50 out of 426 lncRNA-encoded peptides were identified in the seed total proteome during germination (Supplemental table 13), providing strong evidence that the ORFs within these lncRNA are translated and may play a functional role during seed germination.

### uORFs impact the translation of main ORFs during seed germination

uORFs are known to regulate the translation of their downstream main ORF (mORF) by interrupting ribosome scanning during translation (Meijer and Thomas, 2003; Yamashita et al., 2017). To evaluate whether uORFs influence mORF translation during seed germination, the fold change in RF read counts for uORFs between the two consecutive time points during germination was compared to that of the corresponding mORFs (Figure 3a-d). Globally, RF changes at uORFs agree with the changes at the mORF, suggesting that the majority of genes are not translationally regulated by their uORFs. However, comparison across consecutive physiological stages identified nine genes (FDR < 0.05) that exhibit a negative correlation between uORF and mORF translation, characterized by either an increase in RF at the uORF and a decrease at the mORF or *vice versa*. These uORF regulated genes were *ALPHA/BETA-HYDROLASES SUPERFAMILY PROTEIN*, *ABH SUPERFAMILY PROTEIN* (AT5G36210), *DE-ETIOLATED 3*, *DET3* (AT1G12840), *PLASMODESMATA CALLOSE-BINDING PROTEIN 5* (*PDCB5*, AT3G58100), *NAD(P)H DEHYDROGENASE B1* (*NDB1*, AT4G28220), *CHLOROPLAST RNA SPLICING2-ASSOCIATED FACTOR 2* (*CAF2*, AT1G23400), *TETRATRICOPEPTIDE REPEAT (TPR)-like superfamily protein* (AT1G07590) and *PYRUVATE ORTHOPHOSPHATE DIKINASE* (AT4G15530) (Figure 3e-h, Supplemental table 14). From the metaprofile, *ALPHA/BETA-HYDROLASES (ABH) SUPERFAMILY PROTEIN* (AT5G36210) uORF is associated with the ribosome in dry seeds while upon imbibition at 6 HAI, the concurrent decrease (*p*-value = 1.24E10^-5^) of uORF associated ribosome and increase (*p*-value = 0.04) of mORF associated ribosome indicates that ribosome was likely stalled in the dry seeds (Figure 3e, Supplemental table 13). In contrast, DET3 (AT1G12840) which substantially increases (*p*-value = 4.48 E10^-5^) in uORF RF upon imbibition at 6 HAI, leading to the decrease (*p*-value = 1.08E10^-4^) of its main ORF RF (Figure 3e). However, upon further imbibition at 26 HAI, the uORF associated RFs start to decrease, corresponding to an increase of its mORF (Figure 3f), exhibiting a development dependent uORF mediated translation regulation of its downstream genes. The trade-off between the RFs on uORF and mORF upon germination was also seen in other stages at 26-48 HAI and 48-72 HAI (Figure 3g and f). The genes described are representatives of possible uORF mediated translational control related to plant development during seed germination, of which the molecular mechanism can be further explored.

**Figure 3.**
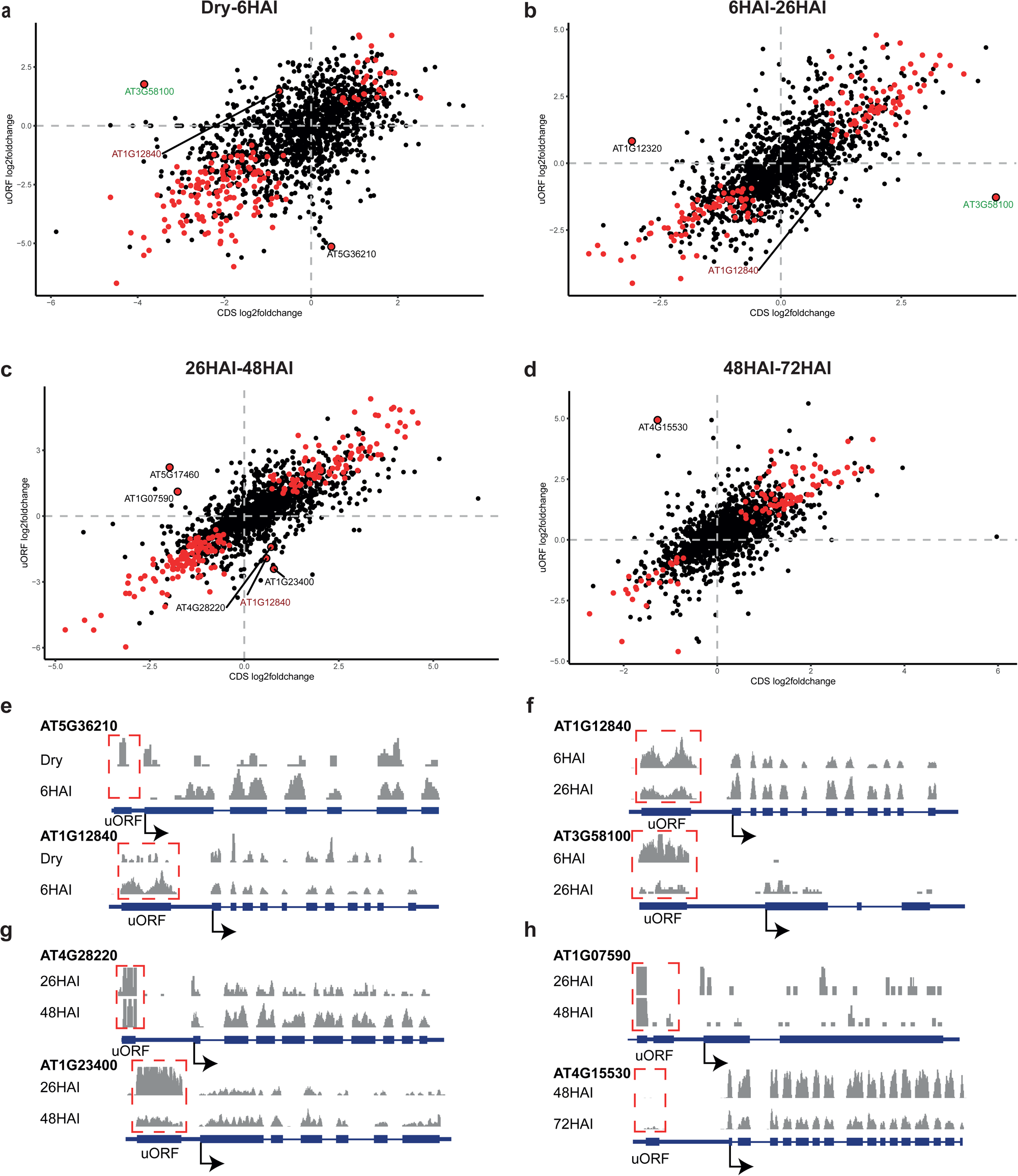
uORF related translational changes during seed germination. (a-d) Scatter plots show the log_2_ fold changes in ribosome footprints for the CDS versus the uORF region for ribosome-associated transcripts across four germination transitions: dry to 6 HAI (a), 6-26 HAI (b), 26-48 HAI (c) and 48-72 HAI (d). Red dots indicate genes with significant changes in ribosome occupancy for the stages that are compared. Genes labelled with AGI codes are those with significant shifts in ribosome distribution during germination progression. AGI labels coloured in red and green represent genes for which the ribosome occupancy changed significantly at multiple transitions. (e-h) RF metaprofiles of genes with either an increase in RF at the uORFs and a decrease at the mORF or *vice versa*. The red frames indicate the uORF region of each gene.

## Discussion

The role of mRNA storage in dry seeds and the regulation of their translation during seed germination is a key question in the field of seed biology. The evidence so far that translation of these stored mRNAs is essential and sufficient for seed germination lies in the fact that a translation inhibitor such as cycloheximide can completely inhibit germination while a transcription inhibitor cannot (Rajjou et al., 2004). Dry seeds are translationally inert, following water uptake during seed imbibition the translational machineries are quickly activated. This results in an increase in the number of ribosomes that are associated to transcripts during seed germination (Bai et al., 2017). Understanding this pattern of ribosome association can provide mechanistic insight into seed germination control. By deep sequencing ribosome-protected fragments at five distinct physiological stages of seed germination, we generated a detailed ribosome occupancy profile of this developmental process at single codon resolution. In dry seeds, and at 6 and 26 HAI seeds 28-nt sized ribosome footprints are most pronounced. In contrast at later germination stages the 29-nt footprints that have been previously reported for Arabidopsis, become more prevalent (Bazin et al., 2017; Gelsinger et al., 2020; Glaub et al., 2020).

The position of ribosome association on an mRNA transcript is pivotal for its translational outcome, since the regions such as the 5’ UTR, the CDS and the 3’ UTR have distinct roles in regulating translation. In dry seeds, ribosome association is significantly higher in both the 5’UTR and 3’UTR compared to germination stages, suggesting a higher proportion of untranslated mRNAs. This pattern is consistent with the absence of translation in dry seeds (Figure 1b, c). Based on previous reports showing that transcripts in dry seeds are predominantly associated with monosomes, we conclude that a ribosome on a given transcript it typically associated with either the 5’ UTR or 3’ UTR, but not both (Bai et al., 2020). Upon imbibition, the translation of stored mRNAs follows two distinct patterns. Transcripts are either translationally down-regulated (Figure 2a, b) or up-regulated (Figure 2c,d), as indicated by changes in the RF distribution along transcripts (Figure 1a, b, Figure 2). These two translational trajectories in seeds correspond to two hypotheses regarding the fate of seed-stored mRNA respectively. 1) Downregulated upon germination: This group includes mRNAs that are actively translated during seed maturation, but degraded upon germination. Examples include transcripts encoding LEAs and seed oil body associated proteins involved in seed filling. This pattern likely represents the gene expression program during seed maturation, where mRNA translation is gradually attenuated towards the end of development, resulting in the complete loss of translational activity in dry seeds (Bai et al., 2023). 2) Upregulation upon germinations: These are mRNAs that encode proteins required during early seed germination, such as seed storage proteins. These transcripts accumulate during seed maturation (Baud et al., 2008) and have been associated with the establishment of seed desiccation tolerance (Terrasson et al., 2013). Upon imbibition, they become translationally active which is in agreement with the observation that storage proteins are synthesized during early germination (Rajjou et al., 2004; Galland et al., 2014). The translation of seed stored mRNAs at early stages of seed imbibition might have ecological advantages. Early imbibition reflects a period in which seeds can still tolerate desiccation (Maia et al., 2011), therefore the translation of these seed maturation and stress response proteins could help tolerating recurrent hydration-dehydration cycles. Moreover, seed storage proteins have been reported as “buffering proteins” for protection against oxidative damage (Nguyen et al., 2015). Their translation might be required for maintaining seed vigour in adverse environments, to increase to chance of seedling survival. This stress surveillance mechanism is specific for seeds before testa rupture and coincides with the stage at which seeds do not tolerate desiccation anymore (Maia et al., 2011). After this stage the transcript abundance and translation of seed storage proteins are sharply reduced (Figure 2c).

Ribosome stalling during protein translation has significant biological roles, for instance to provide enough time for molecular processes like nascent peptide folding, and the recruitment of signal recognition particles for ER targeting (Tanner et al., 2009; Nyathi et al., 2013; Bui and Hoang, 2016; Mercier et al., 2017). The amino acids glycine, aspartate, tyrosine and proline provide the codons for ribosome pausing during seed germination (Figure 1c). This is consistent with previous observations showing that proline, glycine and aspartate are the most conserved amino acids associated with ribosome stalling in yeast, fruit fly, zebrafish, mouse and human (Chyzynska et al., 2021). Ribosome stalling also frequently occurs at aromatic amino acids tyrosine (also identified during seed germination, figure 1c), phenylalanine and tryptophan in bacteria (Cymer et al., 2015). The detection of the context sensitive Kozak consensus sequence that contributes to the ribosome pausing at the start codon in dry seeds suggests a potential sequence-based signal for ribosome stalling (Figure 2d). Although the increased frequency of A residues at -2 and -3 position of Kozak consensus sequence in these seed stored mRNAs may explain ribosome pausing at the start codon, the mechanism and functional relevance of this pausing specifically for these transcripts remains to be discovered. It has been shown that A residues at position -1 to -5 are required for high-level translational efficiency (Kim et al., 2014). Therefore, the increased frequency of A residues at -2 and -3 might contribute to the ribosome pausing in these transcripts. In addition, the high frequency of the G residue at the +4 position constrains the choice of the codon following the start codon, favouring codons beginning with G. This constraint may further increase selectivity for the peptides encoded by these mRNAs.

The identified Kozak consensus sequence in start codon enriched transcripts likely mediates a Kozak motif-dependent translational control in seeds. This is consistent with the strong ribosome peak observed at start codon in dry seeds, which sharply reduces upon water uptake (Figure 1b). Moreover, these findings align with the high polysome association of these transcripts during seed maturation and their reduced association during early germination (Figure S4). Kozak motif-dependent translational control has also been reported for photosynthesis-associated nuclear genes (Hang et al., 2024), suggesting that this mechanism may play a broader role in regulating translation in plants.

lncRNAs have been historically regarded as non-coding RNA based on their size and coding potential. However, there is increasing evidence that lncRNAs can encode short peptides that play roles in transcriptional regulation, splicing and protein scaffolding (Bazin et al., 2017; Zeng et al., 2018; Sebastian-delaCruz et al., 2021; Wu et al., 2024). Moreover, the lncRNA encoded small peptide microRPG1 plays a key role in controlling maize seed desiccation through mediating ethylene signalling (Yu et al., 2024). In this study, we have identified over a thousand of lncRNA that are associated with ribosomes (Supplemental table 11). The identification of peptides that are encoded by lncRNA ORFs provides strong evidence that these lncRNAs are translated during seed germination (Supplemental table 13), which might add an additional layer of gene expression control during seed germination. Not all ribosome-associated lncRNAs contain an ORF which could allow them to generate peptides, however for these lncRNAs a regulatory role during translation cannot be excluded. Although largely unannotated, lncRNAs have been reported to play a role in regulating seed germination, with examples like as *asDOG1* and TE-lincRNA11195 (Wang et al., 2017; Yatusevich et al., 2017). These unexplored resources of ribosome associated lncRNA opens up a plethora of possibilities to understand the translational regulation in seeds.

Translation can also be regulated by uORFs. Well-studied examples in plants are the Conserved Peptide uORF (CPuORF) of the group S basic region leucine zippers (S1 bZIPs), which mediate Sucrose-Induced Repression of Translation (SIRT). SIRT confers the gene expression regulation of the S1 bZIP ATB2/AtbZIP11 upon sucrose treatment (Rook et al., 1998; Wiese et al., 2004; van der Horst et al., 2020). The interaction between uORF-encoded peptides and the ribosome exit tunnel, where the nascent peptide is released on the ribosome, has been shown to regulate the ribosomal arresting, which causes the ribosome stalling (Yamashita et al., 2017). In this study, uORFs that regulate gene expression at the translational level were identified. For example, the translation of the uORF of ALPHA/BETA-HYDROLASES (ABH) SUPERFAMILY PROTEIN gene (AT5G36210) in dry seeds represses the translation of the ABH mORF, whereas at 6 HAI the repression is attenuated (Figure 3e, Supplemental table 13). The function of the ABH family protein during germination remains unknown, however the ABH domain can serve as a core component in the perception of gibberellin, strigolactone and karrikin, which are known to promote seed germination (Shimada et al., 2008; Guo et al., 2013; Hernandez-Garcia et al., 2021). This uORF-mediated gene expression attenuation can also occur upon imbibition, as observed for DET3 (AT1G12840) where increased ribosome association with the uORF correlates with reduced ribosome association with the mORF from dry to imbibed seeds (Figure 3a, e). This uORF-mediated repression is specific to 6 HAI, by 26 HAI the ribosome association with the uORF reduces, coinciding with an increase in mORF ribosome association (Figure 3b, c, f). Development dependent uORF control of gene expression is seen in all stages during germination, which fine-tunes gene expression regardless of their transcript level. uORF mediated translational control has been described as key in fast adaptive response to environmental stress (Chen and Tarn, 2019). Instant gene expression control during seed imbibition is essential to regulate germination, ensuring it only occurs under conditions that are optimal for seedling establishment.

In conclusion, this study provides the first translational landscape at sub-codon resolution across seed germination and identified different mechanisms that may regulate translation during seed germination. The RFs in all germination stages showed *bona fide* ribosome association and substantial ribosome dynamics. In dry seeds, we found relative high ribosome association with the 5’ and 3’ UTRs, and low association with the CDS, compared to later stages of seed germination where association predominates in the CDS. This finding corresponds to the changes in translational status during seed germination. Translation of lncRNAs is supported by their ribosome association as well as the identification of their encoded peptides. The function of these peptides in seed germination has yet to be elucidated. Finally, novel uORFs, that might adjust translation of the mORF, were identified. We propose that the various mechanisms of translational control during seed germination are crucial for enabling rapid responses to environmental changes, thereby ensuring the survival of the germinating seeds and contributing to the seed’s role in the plant’s overall survival strategy.

## Acknowledgements

This work is part of the research domain Applied and Engineering Sciences, project number 15228, which is financed by the Dutch Research Council (NWO). We thank Johannes Hanson and Amir Mahboubi for fruitful discussions during the Ribo-Seq optimalisation and Sjors van der Horst for critically reading the manuscript.

## Author contributions

B.B. and L.B. designed the experiments. B.B. performed the experiments. B.B., H.N. and L.B. directed the design of analysis approaches. B.B., R.Q. and W.S. conducted the analyses. B.B. and L.B. drafted the manuscript. All authors participated in revising and editing the manuscript.

## Figures legends

**Figure S1.**
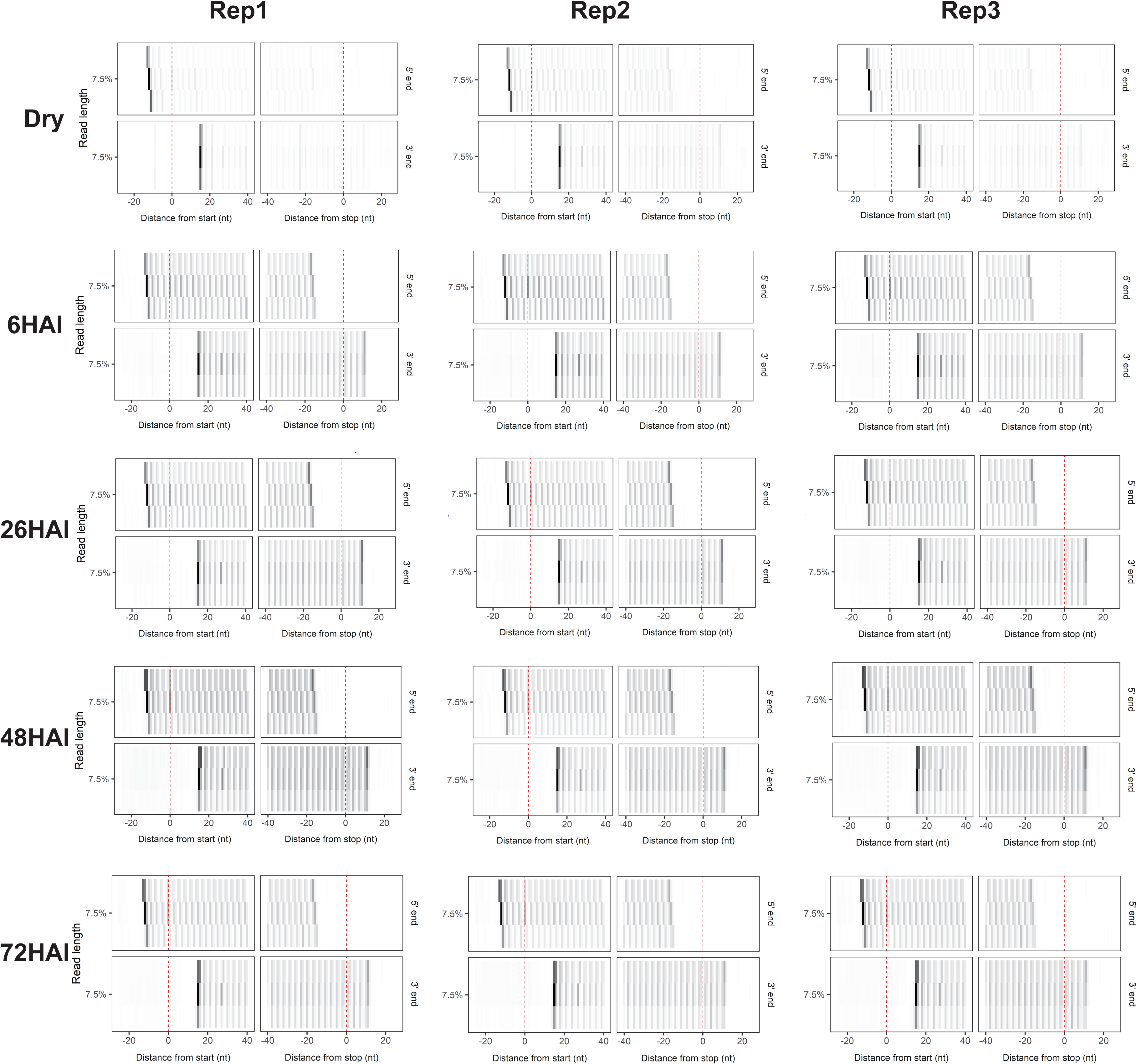
Meta-gene heatmap displaying the signal associated with the 5’ end (upper panel) and 3’ end (lower panel) of the ribosome footprints around the start and the stop codon for fragment lengths 27, 28, 29 nt following dry, 6, 26, 48 and 72 hours after imbibition (HAI) with all three biological replicates.

**Figure S2.**
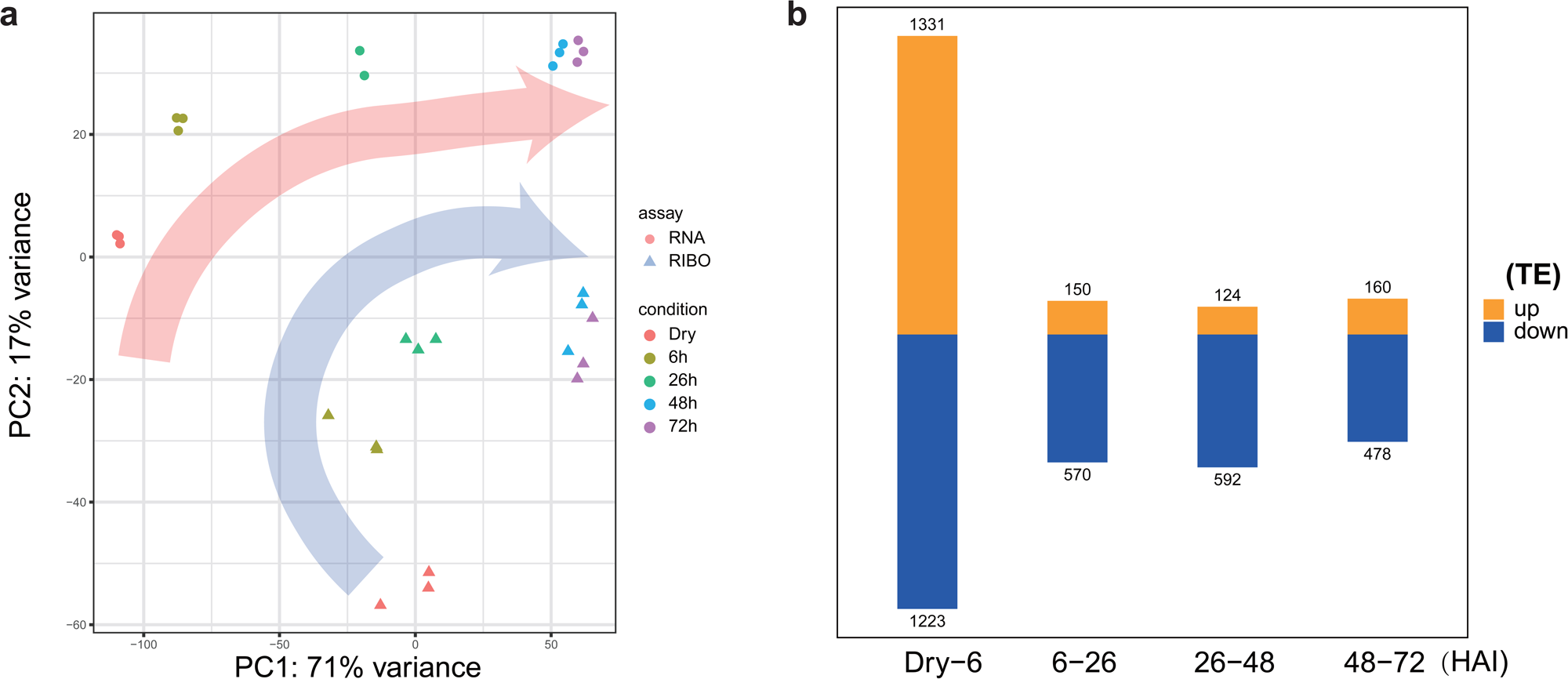
Translational regulated genes during seed germination. (a) PCA plot of total RNA-seq and Ribo-seq data during seed germination. (b) Number of genes with a changed translational efficiency (TE) for different transitions during seed germination.

**Figure S3.**
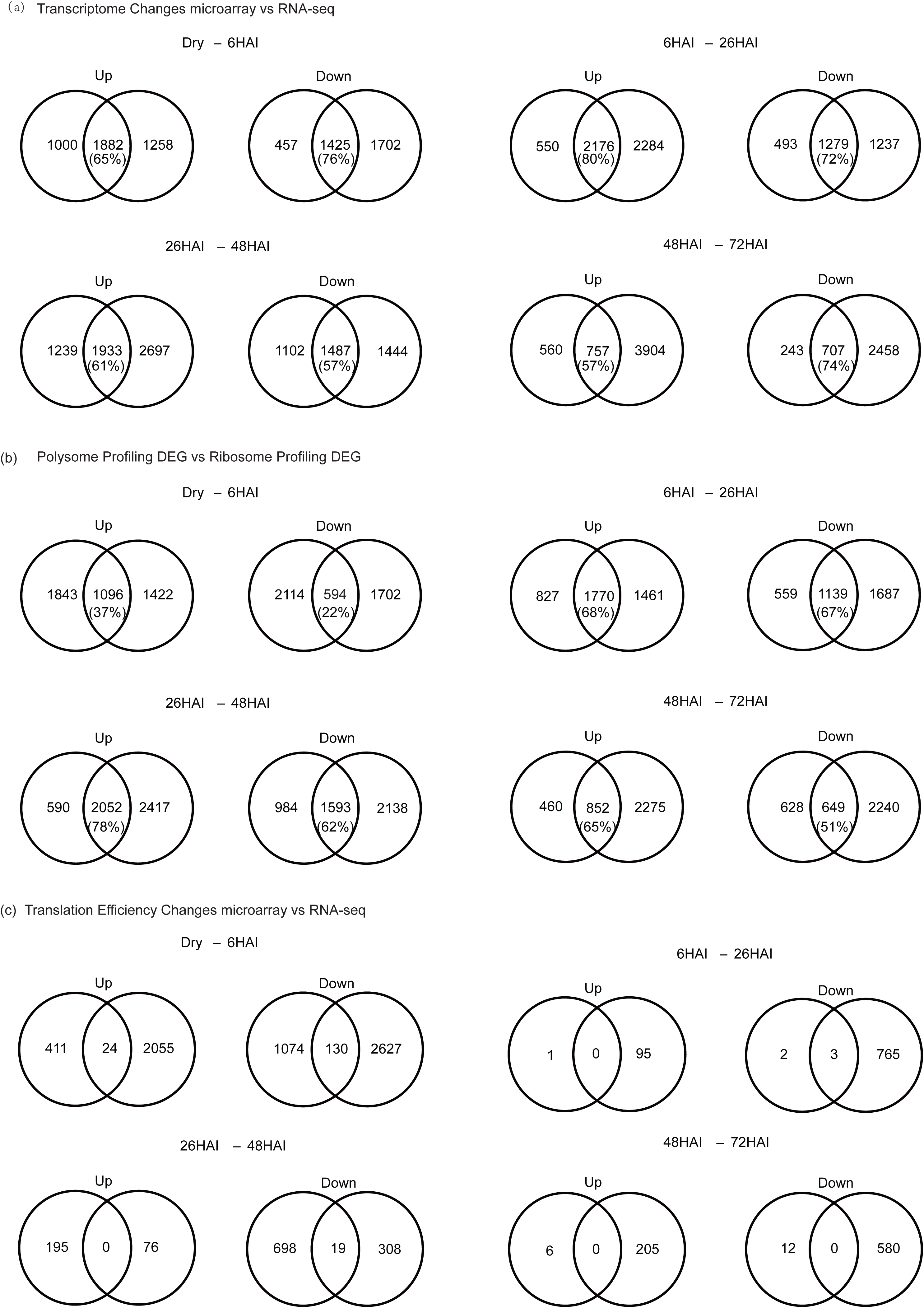
Comparison between results obtained using ribosome sequencing in this study and the polysome profiling analyses during seed germination (Bai et al., 2017) at the (a) transcriptome level, (b) translatome level and (c) translation efficiency level.

**Figure S4.**
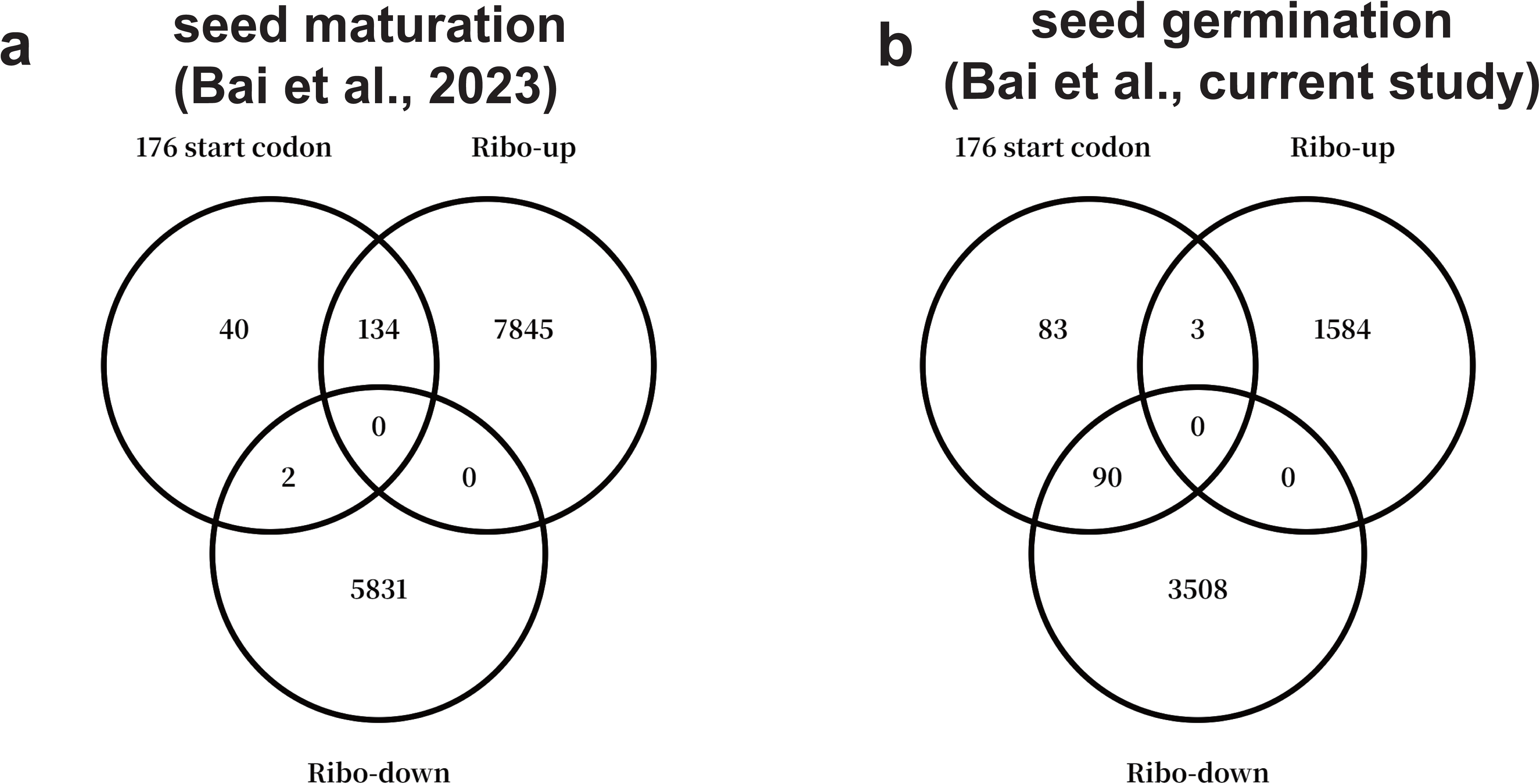
Investigation of the ribosome association of the 176 dry seeds start codon enriched genes during seed maturation and seed germination. (a) Venn diagram representing the overlap of the dry seed start codon enriched genes with polysome associated transcripts during seed maturation. Polysome-up and polysome-down refer to the sum of transcripts that show increased or decreased polysome association during seed maturation as retrieved from (Bai et al, 2023) (b). Venn diagram representing the overlap of the start codon enriched genes with ribosome associated transcripts during seed germination that are retrieved in the current study. The transcripts indicated as Ribo-up or Ribo-down represent the sum of transcripts that have an increased or decreased ribosome association respectively.

## Notes

### Competing Interest Statement

The authors have declared no competing interest.

